# Impact of dupilumab and upadacitinib treatment on epigenetic and transcriptomic alterations in skin-homing T cells of atopic dermatitis patients

**DOI:** 10.1101/2025.04.28.650924

**Authors:** Ahmed MI Elfiky, Hidde M. Smits, Marlot van der Wal, Coco Dekkers, Celeste M Boesjes, Julia Drylewicz, Marjolein de Bruin-Weller, Femke van Wijk

## Abstract

**Background:** Biologic therapies, such as the IL-4 receptor alpha blocker dupilumab, and Janus kinase (JAK) inhibitors like upadacitinib, have revolutionized the management of atopic dermatitis (AD). However, prolonged treatment is necessary to sustain disease remission. While both therapies are clinically effective, only dupilumab has demonstrated successful dose reduction, suggesting potential disease-modifying effects. This study investigates the immune-modulating potential of one year treatment with dupilumab or upadacitinib in AD by assessing epigenetic and transcriptomic changes in skin-homing T cells.

**Methods:** Using flow cytometry, CD4^+^CLA^+^ T cells were isolated from AD patients (n=12) at baseline and after 52 weeks of dupilumab (n=6) or upadacitinib (n=6) treatment. DNA methylome, RNA transcriptome and cytokines expression analyses were conducted using the EPIC array, RNA sequencing and flow cytometry, respectively. Non-atopic healthy controls (HC) (n=6) were included for comparison.

**Results:** We identified 747 differentially methylated regions (DMRs) between HC and AD patients (p-value < 0.1), with a predominance of hypomethylation in AD. These DMR-associated genes were enriched in pathways such as cytokine-cytokine receptor interactions and JAK-STAT signaling. Several DMRs were associated with corresponding changes in the transcriptome and significant differential methylation was observed near 11 AD-associated SNPs. Following treatment, dupilumab partially corrected 6 AD-related DMRs towards the HC profile. Conversely, upadacitinib modulated 40 AD-related DMRs, most shifting further towards the AD profile. Intriguingly, 241 DMRs (upadacitinib) and 13 DMRs (dupilumab) were altered independently of AD-related DMRs, suggesting a broader epigenetic impact of upadacitinib.

**Conclusion:** Our findings demonstrate distinct epigenetic effects of dupilumab and upadacitinib in skin-homing T cells from AD patients. The direction and extent of these modifications may contribute to differences in long-term disease control and have clinical implications for therapy discontinuation strategies.

**Key Messages:**

- Skin-homing T helper cells show unique DNA methylome in AD.
- Dupilumab induces targeted DNA methylome changes, partially rectifying some AD-related DMRs.
- Upadacitinib triggers broader DNA methylome changes, augmenting AD-related DMRs and inducing new DMRs.

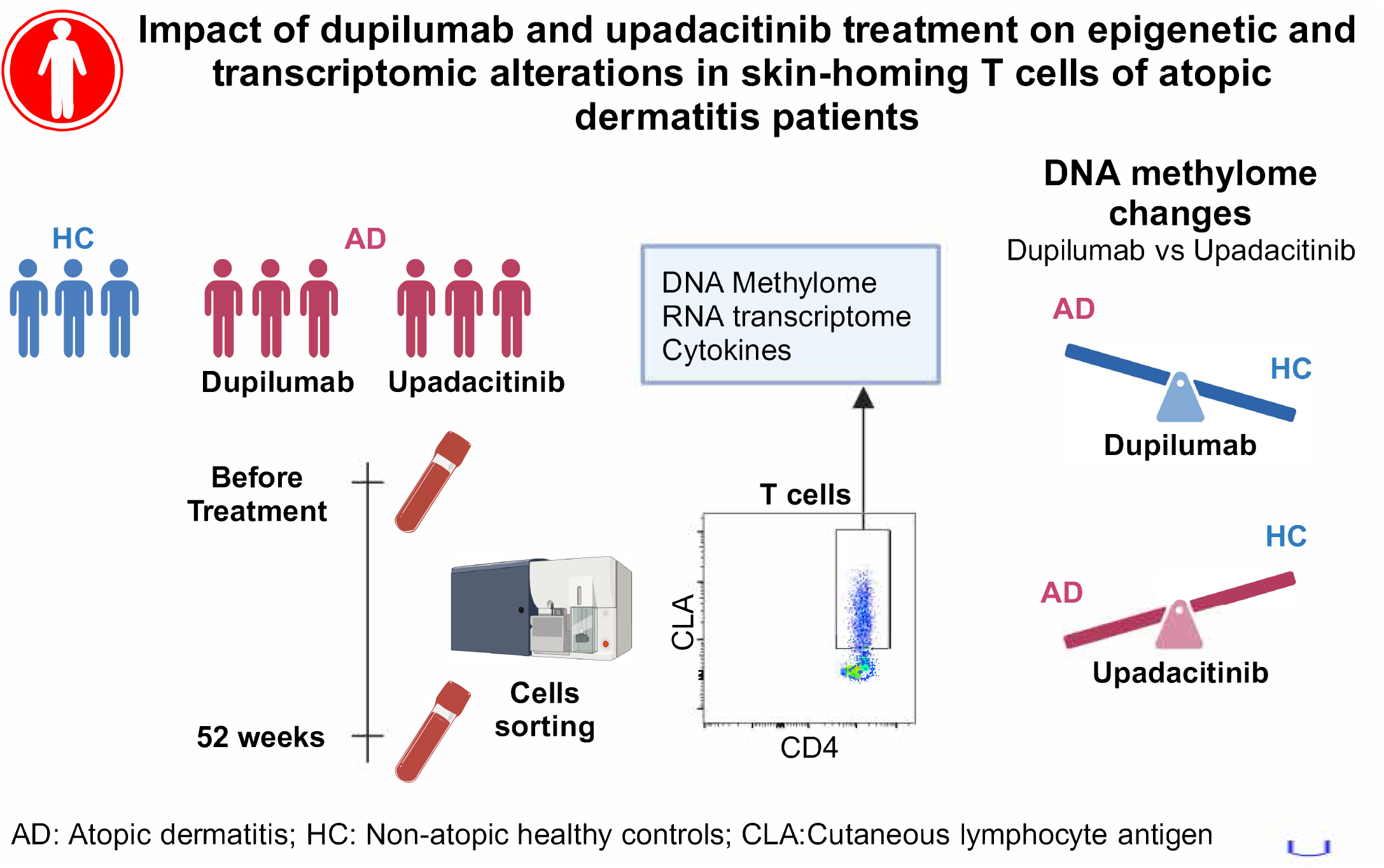

## Introduction

Atopic dermatitis (AD) is a chronic relapsing and remitting inflammatory skin disease that is characterized by skewed T-helper 2 (Th2) mediated immune responses with a predominance of IL4, IL5, IL13 and IL22 cytokines [1]. New biologics and small molecules have been introduced in the last few years, that target specific aspects of AD pathogenesis and are now commonly prescribed in AD patients [2]. These include dupilumab, an IL4 receptor alpha blocker, and upadacitinib, a selective janus kinase (JAK) 1 inhibitor. Randomized clinical trials and real-life data demonstrate excellent clinical efficacy and good safety profile for both medications [3]. In order to prevent disease flares, patients are required to use maintenance therapy for long periods. However, recent clinical studies have shown successful dose reduction of dupilumab while retaining AD remission [4, 5]. Moreover, a recent meta-analysis reported less occurrence of co-morbid allergic events in patients treated with dupilumab [6], indicating an effect that extends beyond AD to other atopic comorbidities. Following treatment discontinuation, dupilumab is associated with longer duration of disease remission [7] compared to upadacitinib, [8]. Whether this collectively reflects a superior disease modifying property of dupilumab is unknown [9].

DNA methylation is an epigenetic mechanism that can alter gene expression without affecting DNA sequence and shape the cellular phenotype and functions in a cell-specific manner. Hyper-or hypo-methylation of CpG islands within a gene promoter or enhancer is associated with reduced or increased gene expression, respectively [10]. In the context of AD, DNA methylation studies have identified an aberrant DNA methylome in lesional skin [11] as well as in peripheral skin-homing CD4^+^CLA^+^ T cells [12]. Skin-homing CD4^+^CLA^+^ T cells, which are recruited into inflamed skin lesions, play an important role in the local inflammatory process, and reflect systemic and local immune alterations under treatment [13]. It is largely unknown whether long-term treatment of current therapies may rectify AD-related epigenetic profiles or induce new changes relevant to disease modification. In the current study, we defined AD-related DNA methylome and RNA transcriptome changes in skin-homing CD4^+^CLA^+^T cells and investigated changes induced by one year treatment with dupilumab or upadacitinib in AD patients.

## Methods

### Patient inclusion

This study included AD patients who presented at University Medical Center of Utrecht (UMCU) dermatology outpatients clinics, as a part of a larger prospective, observational cohort (the Dutch BioDay registry, ClinicalTrials.gov identifier: NCT03549416). Adult patients with moderate-to-severe AD **(Table 1)** were treated with dupilumab or upadacitinib in daily practice and blood samples were collected from patients before initiation of treatment (baseline) and after 52 weeks of treatment. All patients signed institutional review board–approved written, informed consent, adhering to the Declaration of Helsinki Principles.

### Skin-homing T cell sorting and flow cytometry analysis

PBMCs were separated via Ficoll-Paque (GE Healthcare, Chicago, IL) density gradient centrifugation. Following isolation, PBMCs were cryopreserved in RPMI 1640 medium supplemented with 2mM L-glutamine, 100 IU/ml penicillin-streptomycin, 20% fetal bovine serum (FBS), and 10% DMSO (Sigma-Aldrich, St. Louis, MO), then stored at −170 °C until required for further analysis. PBMCs were thawed, counted and 1-2 × 10^6^ cells were used for flow cytometry staining; 2.5-5 × 10^5^ cells were seeded per well for each staining panel. The remaining PBMCs (6-10 × 10^6^ cells) were used for CD3^+^CD4^+^CLA^+^ T cell flow cytometry sorting. Antibodies details are summarized in **Table S1**.

For cells sorting; cells were washed with phosphate-buffered saline (PBS, Sigma-Aldrich), then incubated with eBioscience Fixable Viability Dye (Invitrogen) for 30 minutes at 4°C. Cells were then washed, and incubated with a surface antibody mix **(Table S1)** in FACS buffer for 25 minutes at 4°C, then washed and resuspended in FACS buffer (PBS, 2% FBS, 0.1% sodium azide). Following that, LD^−^CD19^−^CD3^+^CD4^+^CLA^+^ T cells **(Figure S1)** were sorted (50,000 cells) on ARIA-III sorter. Sorted cells are centrifuged and lysed in RLT plus lysis buffer (AllPrep DNA/RNA Micro Kit; Qiagen). Samples were stored in −80C for future analysis.

For intracellular cytokine staining, cells were first restimulated with 20 ng/ml phorbol 12-myristate 13-acetate (PMA; Sigma-Aldrich) and 1.0 μg/ml ionomycin (Sigma-Aldrich) for 4 hours at 37°C in a 5% CO2 atmosphere. Golgistop (0.26% monensin; BD Biosciences, 1:1,500) was added during the last 3.5 hours of stimulation. Following stimulation, cells were collected, washed with PBS and incubated with eBioscience Fixable Viability Dye (Invitrogen) for 30 minutes at 4°C. Cells were then washed, and incubated with a surface antibody mix **(Table S1)** in FACS buffer for 25 minutes at 4°C. Cells were fixed and permeabilized sequentially with eBioscience fixation/permeabilization reagent (Invitrogen) at 4°C for 30 minutes each step. Cells are then washed and incubated with an intracellular antibody mix **(Table S1)** at 4°C for 30 minutes. Samples were then analyzed on an LSR Fortessa either on the same day or the next day.

### Nucleic acid isolation, RNA-sequencing, and DNA methylation EPIC array

Skin-homing T cells were collected and stored in RLT plus lysis buffer. They were thawed on ice, followed by simultaneous isolation of RNA and DNA using AllPrep DNA/RNA Micro Kit (Qiagen) and following the manufacturer’s protocol. The quality of isolated DNA and RNA was checked on tapestation, using genomic DNA screentape analysis kit (Agilent) and high sensitivity RNA screentape analysis kit (Agilent) respectively. Samples with DIN score > 7 for DNA and RIN score > 7 for RNA were sent for subsequent analysis. DNA methylation analysis using Illumina’s Infinium MethylationEPIC v2.0 935k BeadChip and next generation RNA sequencing analysis were performed by GenomeScan B.V. (Plesmanlaan 1d, 2333 BZ, Leiden, The Netherlands).

#### For RNA sequencing

The NEBNext Ultra II Directional RNA Library Prep Kit for Illumina was used to process the samples. The sample preparation was performed according to the protocol “NEBNext Ultra II Directional RNA Library Prep Kit for Illumina” (NEB #E7760S/L). Briefly, mRNA was isolated from total RNA using the oligo-dT magnetic beads. After fragmentation of the mRNA, a cDNA synthesis was performed. This was used for ligation with the sequencing adapters and PCR amplification of the resulting product. The quality and yield after sample preparation was measured with the Fragment Analyzer. The size of the resulting products was consistent with the expected size distribution (a broad peak between 300-500 bp). Clustering and DNA sequencing using the NovaSeq6000 was performed according to manufacturer’s protocols. A concentration of 0.8 nM of DNA was used. NovaSeq control software NCS v1.8 was used. Primary data analysis and results Image analysis, base calling, and quality check were performed with the Illumina data analysis pipeline RTA3.4.4 and BclConvert v4.2.4. Data is delivered in fastq.gz format.

#### For DNA Methylation

Genomic DNA was bisulfite-converted using the EZ DNA Methylation Gold Kit (Zymo Research) and used for microarray-based DNA methylation analysis, performed at GenomeScan (Leiden, The Netherlands) and AIT (Austrian Institute of Technology GmbH, Vienna, Austria) on the Human Methylation 935k BeadChip (Illumina, Inc., San Diego, CA, USA). The bisulfite converted DNA was processed and hybridized to the HumanMethylation EPICv2 BeadChip (Illumina, Inc.), according to the manufacturer’s instructions. The BeadChip images were scanned on the iScan system, and the data quality was assessed using the R script MethylAid (M. van Iterson, 2023) using default analysis settings. All samples showed a detected CpG (p=0.01) above 96%, > 897k detected CpG (p=0.01).

### Bioinformatics analysis

#### DNA Methylation analysis

The DNA methylation data used in this study were generated from Illumina Infinium Human Methylation BeadChip v2.0 arrays, with raw intensity data provided in .idat files. Raw signal intensities were processed with the R package minfi (version 1.44.0) [14]. Quality control assessment to ensure data integrity was made by checking for sample outliers and gender mismatches. All samples passed this step. Low-performing probes were filtered out, after which data were preprocessed and normalized with the preprocessFunnorm function from minfi. The resulting data were transformed to M-values, with probes on sex chromosomes, probes known to cross-hybridize, and SNP-associated probes (MAF > 0.05) were removed using rmSNPandCH from the package DMRcate (version 1.8.6) [15]. M-values for probes matching the same genomic site were averaged.

M-values were used in subsequent statistical analyses to assess associations between DNA methylation, disease, and treatment. A UMAP of the baseline samples was generated using the package uwot (version 0.2.2) [16] with the following parameters: n_neighbors = 3, min_dist = 0.05. Differentially methylated positions (DMPs) were identified with the limma package (version 3.60.4) [17], using age and gender as covariates. For the unpaired analyses (e.g., HC vs. AD samples), DMRs were identified using the package CoMethDMR (version 3.20) [18]. Genic regions were defined based on the UCSC hg38 gene annotation, and intergenic regions had a minimum of three probes within 200 bp. Gene set enrichment analysis was conducted using a functional enrichment test (FET) from the clusterProfiler package (version 4.14.6) [19], focusing on genic DMRs and leveraging the KEGG pathway database for annotation.

For the subsequent analyses, comparing HC against AD samples and AD baseline (BL) against AD samples after 52 weeks of treatment, the package mCSEA (version 1.18.0) [20] was used to increase sensitivity in identifying DMRs. Custom promoter annotations were created using the EPIC-8v2-0_A1.csv file from Illumina, with probes mapped to the following genomic features; TSS1500, TSS200, 5’UTR, and exon_1. Fold-changes were compared pairwise between BL and 52 weeks of treatment scenario and in an unpaired manner for the HC vs AD. For visualization, mean fold-changes of the leading edge CpGs for significant DMRs were used.

#### RNA sequencing analysis

Paired-end reads were checked for quality using the package FastQC (version 0.11.9) [21] and mapped to the hg38 genome using the package rSubRead (version 2.0.6) [22] with default settings. Read counts were quantified and analyzed in DESeq2 (version 1.38.3) [23]. Analyses between HC and AD samples were unpaired, while those comparing AD baseline and treatment samples were paired. All analyses were conducted in R (version 4.4.1) with a significance threshold set at an FDR of 0.1.

## Results

### Skin-homing T cells exhibit a distinct DNA methylome profile in AD patients

In the current study, we performed combined DNA methylome and RNA transcriptome analysis on flow cytometry-sorted skin-homing T helper cells. CD4^+^CLA^+^ T cells were sorted **(Figure S1)** from PBMCs of non-atopic healthy donors (HC) (n=6) and AD patients (n=12) at baseline and 52 weeks after dupilumab (n=6) or upadacitinib (n=6) treatment, as illustrated in the **graphical abstract**. The median age of HC participants is 28 years old, compared to 29.5 and 26 years in AD patients treated with dupilumab and upadacitinib, respectively. Half of HC participants are male, compared to 66.7% in AD patients groups. All AD patients reported adequate clinical response to both treatments reflected by progressive reductions in severity index (EASI) scores at 4, 16, and 52 weeks of follow-up. It is noteworthy that upadacitinib treated group had a prior exposure to biologic treatment; 5 out of 6 patients were previously treated with dupilumab and 4 out of 6 patients were previously exposed to baricitinib treatment. In contrary, dupilumab treated group was biologic treatment naïve. Both groups had prior exposure to other immunosuppressive medications such as cyclosporin and tacrolimus **(Table 1)**. First, we defined the AD-associated DNA methylome signature in skin-homing T helper cells, by comparing HC to all AD patients at baseline. Uniform Manifold Approximation and Projection (UMAP) dimensional reduction clearly separated the majority of AD samples from HC **(Figure 1A)**. We identified 12,702 differentially methylated positions (DMPs) between HC and AD, comprising 9,489 hypomethylated DMPs and 3,213 hypermethylated DMPs **(Figure 1B)**. This pattern indicates a predominance of DNA hypomethylation in skin-homing T cells of AD patients. The DMPs were distributed across multiple DNA regions, with 1,270 DMPs located in enhancer regions. Additionally, 1,846 DMPs were situated in promoter regions, while 7,922 DMPs were found in gene body regions. Further analysis of the DMPs within promoters and gene bodies, utilizing the CoMethDMR package [20], identified 747 differentially methylated regions (DMRs), comprising 487 hypomethylated DMRs and 260 hypermethylated DMRs **(Figure 1C)**. Among these, 154 DMRs were situated in promoter regions, while 231 DMRs and 265 DMRs were found in gene body and intergenic regions, respectively **(Figure 1D)**. For a comprehensive understanding of genes associated with differentially methylated promoters and enhancers, gene set enrichment analysis (GSEA) was performed. This analysis showed enrichment of various pathways, including cytokine-cytokine receptor interactions and JAK-STAT signaling pathways **(Figure 1E)**. RNA transcription analysis of the sorted skin-homing T helper cells revealed 350 differentially expressed genes (DEGs) between AD patients and HC. Upon overlapping DMRs in promoter regions or DMPs in enhancer regions with DEGs, we identified 48 genes that are modulated in AD at both the methylome and transcriptome levels. Hypomethylation of DNA is generally associated with increased gene expression and vice versa. For 35 out of the 48 genes, changes in methylation were indeed associated with corresponding alterations in gene expression: hypermethylation (8 genes) was linked to decreased expression, while hypomethylation (27 genes) was linked to increased expression. **(Figure 1F)**. Among the DEGs regulated at the DNA methylation level, we identified promoter hypomethylation of gene-encoding *lamin A/C (LMNA*) and *G protein-coupled receptor 55 (GPR55*) in our analysis. It is noteworthy that many DMRs did not overlap with DEGs in our datasets.

**Table 1.** Patients’ characteristics.

| Patients' characteristics | AD dupilumab cohort (n=6) | AD upadacitinib cohort (n=6) |
| --- | --- | --- |
| Age (years), median (IQR) | 29.5 (9) | 26 (14) |
| Gender (Male), n (%) | 4 (66.67%) | 4 (66.67%) |
| AD age of onset, n (%) |  |  |
| - Childhood | 6 (100%) | 5 (83,33%) |
| - Adolescence | - | 1 (16,67%) |
| Atopic comorbidities, n (%) |  |  |
| - Allergic asthma | 3 (50%) | 3 (50%) |
| - Allergic rhinitis | 4 (66,67%) | 3 (50%) |
| - Allergic conjunctivitis | 3 (50 %) | 5 (83,33%) |
| - Food allergy | 2 (33,33%) | 3 (50%) |
| EASI score median (IQR) |  |  |
| - BL | 14.6 (13.6) | 18.9 (13.7) |
| - Week 4 | 10.4 (8.75) | 5.8 (8.3) |
| - Week 16 | 5.1 (1.25) | 5.55 (4) |
| - Week 52 | 2.35 (2.4) | 4.45 (2.7) |
| NRS itch score, median (IQR) |  |  |
| - BL | 8 (3) | 8 (2) |
| - Week 4 | 4 (5.5) | 3 (1) |
| - Week 16 | 2 (3) | 4.5 (3) |
| - Week 52 | 2 (3) | 3.5 (2.7) |
| Treatment adverse events, n (%) |  |  |
| - Conjunctivitis | 5 (83,33%) | - |
| - Infections (herpes zoster) | - | 1 (16,67%) |
| Prior immunosuppressives use, n (%) |  |  |
| - Methotrexate | 2 (33,33%) | 2 (33,33%) |
| - Cyclosporin | 6 (100%) | 5 (83,33%) |
| - Mycophenolic acid | - | 1 (16,33%) |
| Biologics and small molecules, n (%) |  |  |
| - Dupilumab | - | 5 (83,33%) |
| - Baricitinib | - | 4 (66,67%) |

**Figure 1.**
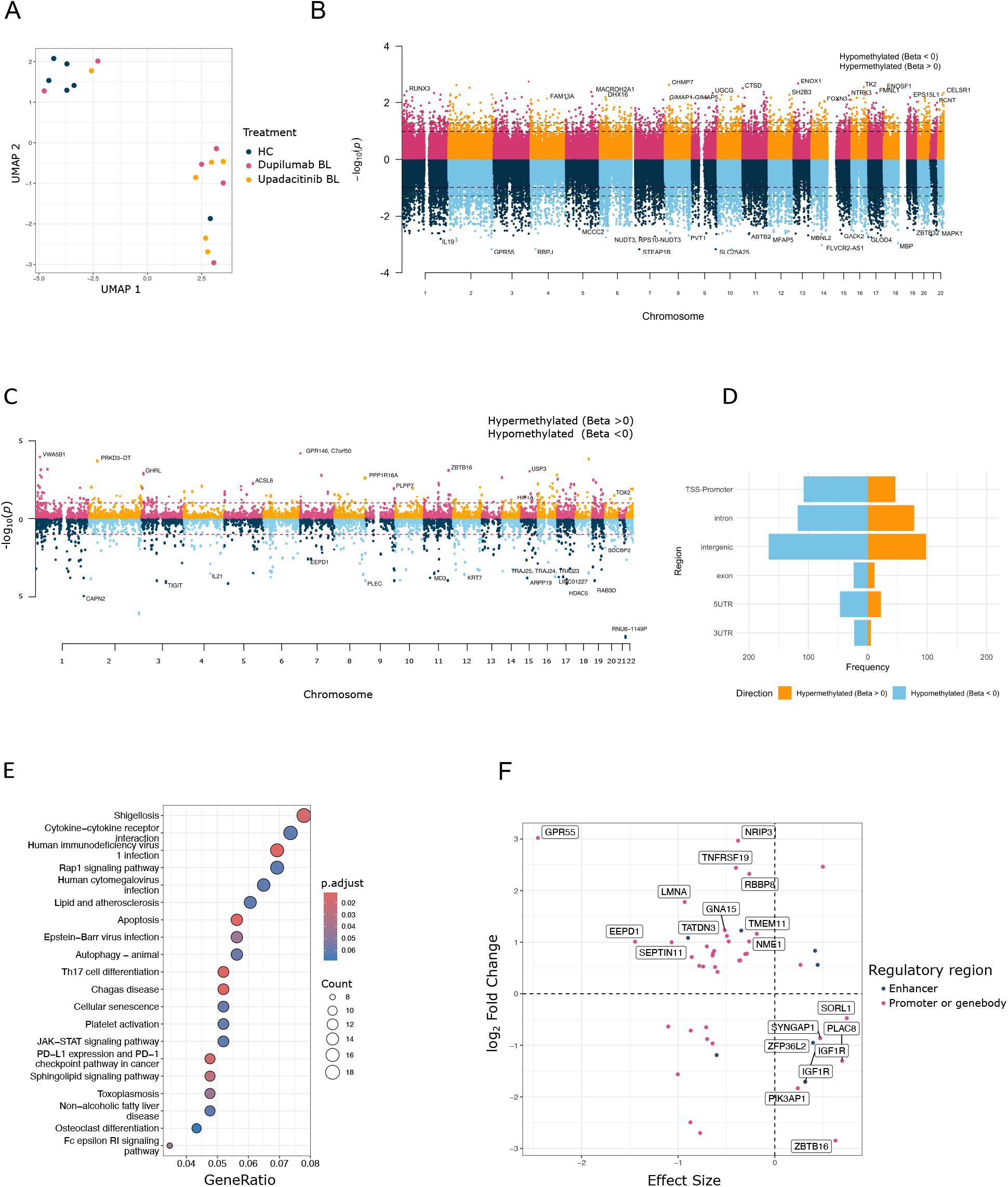
Skin homing T cells exhibit a distinct DNA methylome profile in AD. Comparing atopic dermatitis (AD) patients to healthy non-atopic controls (HC); (A) UMAP visualization of samples from AD patients at baseline (BL; pre-treatment with dupilumab and upadacitinib) and HC. (B) Manhattan plot displaying differentially methylated positions (DMPs) in baseline AD samples compared to HC. Each dot represents a CpG site: hypermethylated CpGs are colored in pink (even chromosomes) and orange (odd chromosomes), while hypomethylated CpGs are navy (even chromosomes) and light blue (odd chromosomes). Genes associated with top DMPs are highlighted. (C) Manhattan plot of differentially methylated regions (DMRs) identified by CoMethDMR in baseline AD samples compared to HC. Regions are represented as dots: hypermethylated regions are shown in pink (even chromosomes) and orange (odd chromosomes), while hypomethylated regions are navy (even chromosomes) and light blue (odd chromosomes). The red dotted line denotes a false discovery rate (FDR) of 0.1 Top DMR-associated genes are highlighted. (D) Bar plot illustrating the genic distribution of hyper-and hypomethylated DMRs. (E) Functional enrichment analysis for genes and promoters within the DMRs identified by CoMethDMR. (F) Correlation between the direction of methylation changes in DMRs (located within a gene, promoter, or within 5 kb of an enhancer) and the log2 fold change in corresponding gene expression levels. Genes exhibiting differences in both gene expression and DNA methylation are labeled.

### Differential methylation pattern between AD and HC is identified near 11 AD-associated SNPs sites

As large genetic analysis studies (GWAS) have previously identified multiple AD-associated single nucleotide polymorphisms (SNPs) [24], we investigated next whether there were changes in the methylation profile at CpG sites located at, or within cis-regulatory distance of these well-characterized AD-associated SNPs. Among 79 known AD-associated SNPs, we identified significant differential methylation within DMPs located within 5KB of 11 SNPs **(Figure 2A)**. We observed significant hypomethylation in close proximity to SNPs associated with *IL13, STAT3, AGO2, CSF2RB* and *TH2LCRR* genes. Additionally, significant hypermethylation was noted at SNPs associated with the *CLEC16A, DOK2, GLB1, IL18RAP, ZNF365*, and *LINC02929* genes **(Figure 2B)**. No significant transcriptomic changes were observed in these genes. However, some genes showed no detectable expression in our dataset.

**Figure 2.**
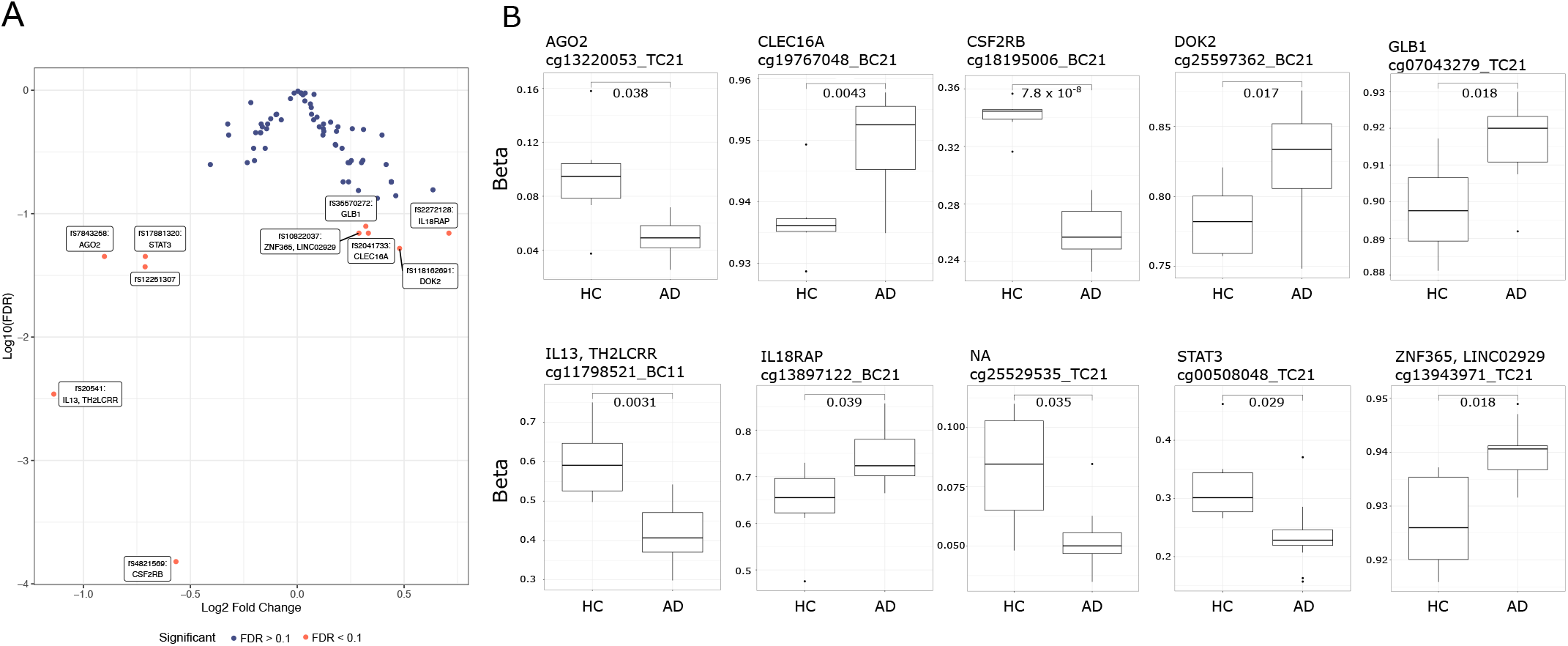
Differential methylation near AD-related SNP sites in AD compared to HC. (A) Volcano plot displaying differential methylation at CpG sites located within 5 KB of AD-associated single nucleotide polymorphisms (SNPs). Significant differentially methylated positions (DMPs) are highlighted and annotated with their corresponding SNPs and associated genes. (B) Boxplots illustrating the beta values (quantification of methylation levels at CpG sites; 0 is totally unmethylated, 1 is totally methylated) of DMPs found near or within SNPs in atopic dermatitis (AD) baseline samples compared to healthy non-atopic controls (HC).

### DNA methylation changes in response to dupilumab or upadacitinib treatment

We next investigated whether dupilumab or upadacitinib could rectify the AD-related DMRs, or induce new, off-target DMRs unrelated to AD. We utilized the mCSEA package to identify altered DMRs within gene promoters, and compared samples taken before and after 52 weeks of treatment with either dupilumab or upadacitinib. We identified 453 differentially methylated AD-related DMRs between AD patients at baseline and healthy controls. When comparing post-treatment to pre-treatment samples, dupilumab modulated 19 DMRs, 6 of which overlapped with previously defined AD-related DMRs. Upadacitinib modulated 281 DMRs, including 40 AD-related regions. In addition to these AD-associated changes, both treatments induced distinct epigenetic alterations. Specifically, dupilumab resulted in the emergence of 13 new DMRs, while upadacitinib led to 241 new DMRs **(Figure 3A)**. Lists of genes annotated to dupilumab or upadacitinib modulated DMRs are provided in **table S2** and **table S3**, respectively. Comparing AD patients treated with dupilumab and upadacitinib at baseline showed no significant differences in the methylation profiles prior to initiation of therapy (data not shown).

**Figure 3.**
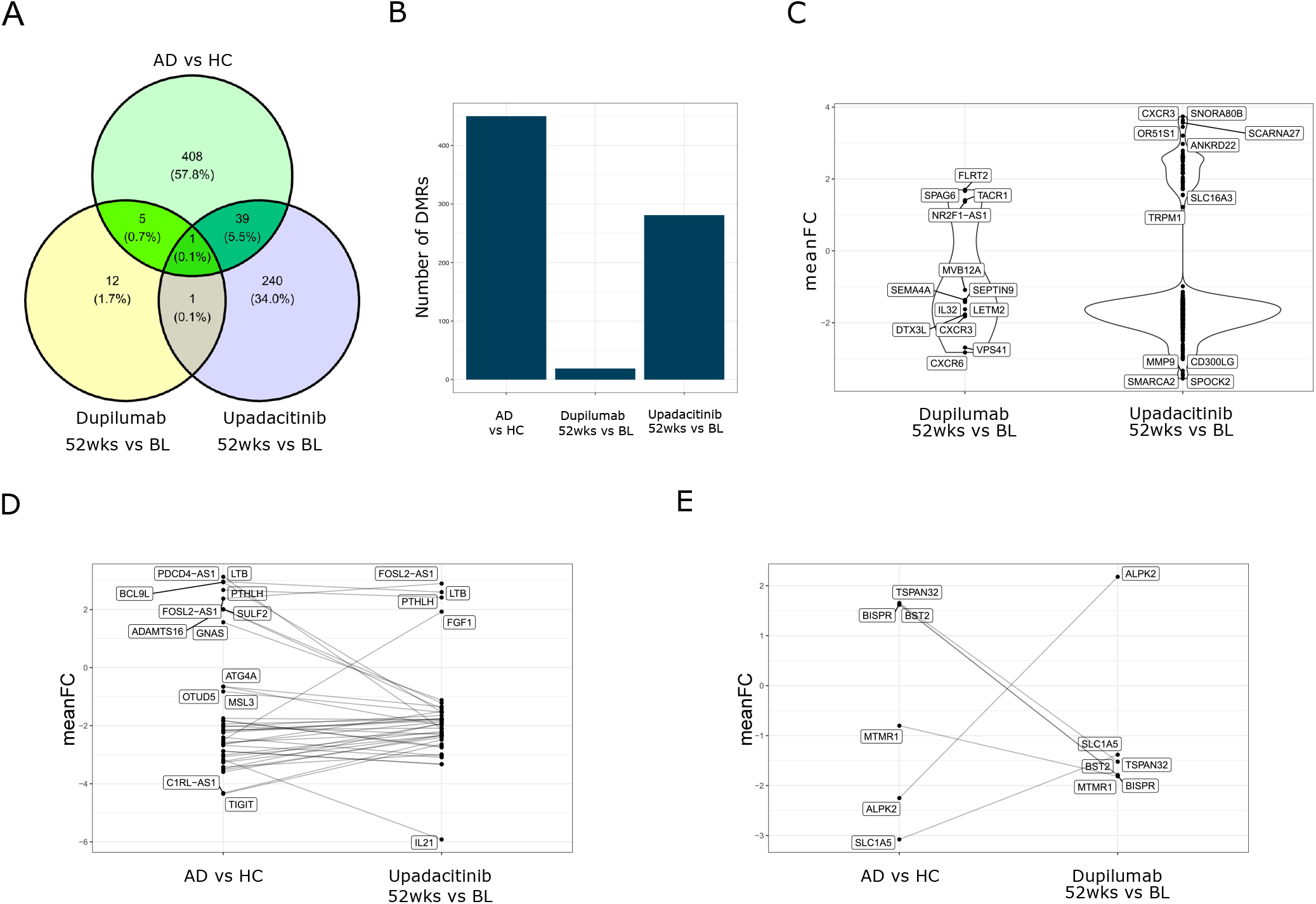
DNA methylation changes in response to dupilumab or upadacitinib treatment. (A) Venn diagram of the differentially methylated promoters (as determined by mCSEA, p value < 0.1) between the three comparisons; atopic dermatitis (AD) patients at baseline compared to healthy non-atopic controls (HC), AD patients after compared to before dupilumab treatment, AD patients after compared to before upadacitinib treatment. (B) Bar plot indicating the total of differentially methylated regions (DMRs) for each comparison. (C) The mean log2 fold change of the leading edge probes for the newly induced differentially methylated genes promoters, in response to dupilumab or upadacitinib treatment. (D) The mean log2 fold change of the leading edge probes for the AD-related differentially methylated genes promoters that change in response to upadacitinib treatment or (E) dupilumab treatment, comparing both AD patients to HC and AD patients after treatment to before treatment.

When comparing the effects of both treatments after 52 weeks, upadacitinib modulated more AD-related DMRs (**Figure 3B**) but also induced a greater number of new DMRs compared to dupilumab **(Figure 3C)**. Interestingly, when we examined the directionality of change, upadacitinib enhanced the majority of AD-related DMRs (34 out of 40), further shifting the methylome profile toward that characteristic of AD **(figure 3D)**. In contrast, dupilumab, reversed 4 out of 6 AD-related DMRs toward a methylome profile more closely resembling HC. The four reversed DMRs are situated within the promoters of *BST2, BISPR, TSPAN32*, and *ALPK2* genes. Among these, all except *ALPK2* exhibited increased methylation in AD patients, which was subsequently reduced after dupilumab treatment. Conversely, *ALPK2* displayed decreased methylation in AD patients, which was corrected after dupilumab treatment **(Figure 3E)**.

### *IL13* and *IL22* are epigenetically controlled at the DNA methylation level, with *IL13* promoter methylation being altered in opposite ways by dupilumab and upadacitinib

AD is a type 2 immune inflammatory disease in which Th2 cytokines play a key role in disease pathogenesis. To investigate the impact of treatment on Th2-driven inflammation, we analyzed both protein expression and promoter methylation status of Th2 cytokine genes in AD, and after 52 weeks of dupilumab or upadacitinib treatment. Flow cytometry revealed an increased percentage of IL4, IL5, IL13, and IL22 protein expressing skin-homing CD4^+^ T cells in AD compared to HC **(Figure 4A)**. At the DNA methylome level, the *IL13* and *IL22* gene promoters, but not the *IL4* or *IL5* gene promoters, were significantly hypomethylated in AD **(Figure 4B)**. No detectable mRNA expression of these cytokines was observed. Interestingly, the *IL13* promoter demonstrated decreased methylation in AD patients that corresponded to elevated IL13 protein expression. Following treatment, dupilumab increased *IL13* promoter methylation levels towards levels observed in HC, whereas upadacitinib further decreased methylation compared to baseline **(Figure 4C)**. Despite these changes, IL13 protein expression remained unchanged following both treatments at week 52 **(Figure 4D)**. Together these findings indicates that IL13 and IL22 expression are epigenetically regulated at the DNA methylome level in skin-homing CD4^+^ T cells and that IL13 is susceptible to modulation by dupilumab or upadacitinib treatment.

**Figure 4.**
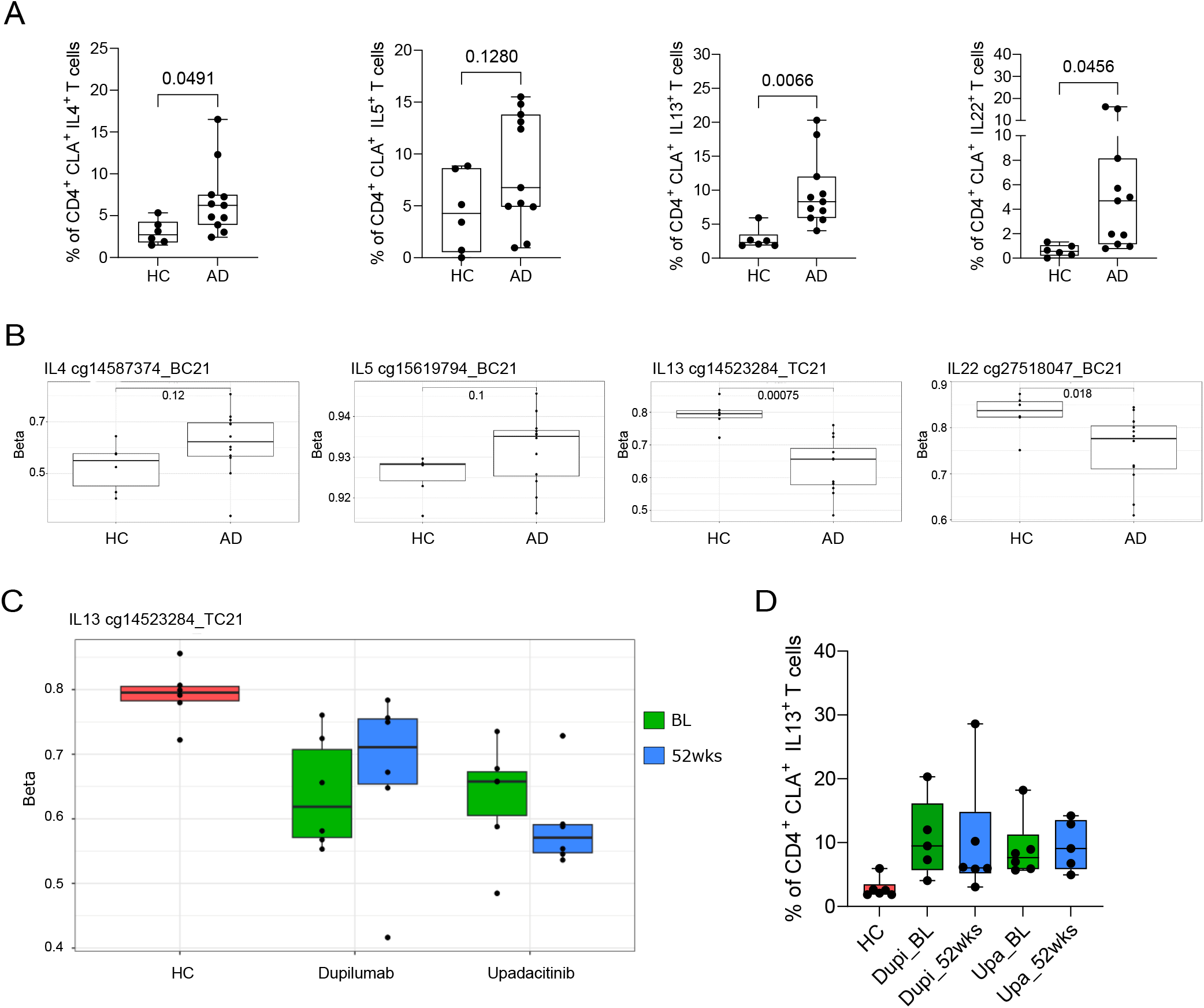
Th2 cytokines protein expression and DNA methylation of gene promoters in AD patients and in response to dupilumab or upadacitinib treatment. (A) frequency of IL4, IL5, IL13 and IL22 expressing skin-homing T helper cells in both healthy non-atopic controls (HC) and atopic dermatitis (AD) patients. (B) Methylation beta value of leading probe within promoters of *IL4, IL5, IL13* and *IL22* genes in both HC and AD patients. (C) changes in methylation beta value of leading probe within promoter of *IL13* following dupilumab or upadacitinib treatment. BL; baseline, 52wks; 52 weeks after treatment, Dupi; dupilumab, Upa; upadacitinib. Student t test is applied, p values are shown.

## Discussion

Biological treatments, such as dupilumab and small-molecules JAK inhibitors like upadacitinib have revolutionized the treatment of moderate to severe AD. However, very little is known about their effect on epigenomic profile of the disease and its potential implications for disease modification. In the current study, we characterized AD-related DNA methylome and RNA transcriptome alterations in skin-homing CD4^+^CLA^+^ T cells. The distinct AD-related DNA methylome is characterized by global hypomethylation, reflecting a predominant gene activation state associated with inflammatory disease. It is noteworthy that many AD-related methylome changes did not correspond to detectable transcriptomic alterations in our datasets. A likely explanation is that inducible genes, including cytokines and chemokines, were not captured in the transcriptomic analysis because skin-homing T helper cells were not stimulated after sorting and prior to RNA sequencing. Consequently, several potentially relevant DMRs were associated with genes exhibiting very low or nearly undetectable expression. Among the interesting genes modulated at both the methylome and transcriptome level, we identified *LMNA, GPR55*, and *Fut7*. LMNA plays a key role in T cell polarization, as *LMNA* deficient T cells demonstrate skewed T cell polarization toward regulatory T cells (Tregs) over Th1 cells [25, 26]. GPR55 is an immune mediator that is involved in immune cell activation. In an AD murine model, GPR55 antagonist ameliorated AD symptoms and reduced T helper cells in regional lymph nodes [27]. *Fut7* encodes an enzyme involved in the generation of selectin ligands, which are crucial for skin-homing properties of T cells [28]. These findings indicate modulation of DNA methylation in genes involved in T cell–mediated inflammation in AD. We further investigated the differential methylation patterns in close proximity to known AD-associated SNPs. SNPs can create or disrupt DNA methylation in the flanking regions upstream or downstream of the genetic variant. This phenomenon is called allele-specific methylation, a pattern of DNA methylation in which differential methylation exists between a pair of alleles, unlike general methylation, which affects both alleles equally. The SNP-created differential methylation pattern contributes to the mechanisms underlying how certain disease associated-SNPs influence disease phenotype [29, 30]. In atopic dermatitis, a previous study showed that the SNP rs7872806 leads to increased methylation at the CpG site cg13920460, impacting *tight junction protein 2 (TJP2)* expression. This, in turn, modulates downstream nuclear protein activator protein 1 (AP-1) and cyclin D1 (CD1) levels, leading to increased cell proliferation and transepidermal water loss, which suggests a role in the development of epidermal hyperplasia [31]. Among previously reported AD-associated SNPs, we observed significant differential methylation in close proximity to 11 SNPs, including differential hypomethylation near rs20541 (*IL13*) and rs17881320 (*STAT3*). This suggests a potential role for methylation changes within close proximity to the 11 AD-associated SNPs in the pathogenesis of the disease. In our study, we did not include SNP genotyping within the same cohort, therefore, we cannot correlate the differential methylation patterns identified near the SNPs sites with their presence. Further studies are needed to investigate whether these 11 SNPs create differential methylation pattern in individuals carrying those SNPs, which, in turn, can influence gene expression. This will provide further mechanistic insight into how these SNPs may influence disease phenotype via epigenetic mechanisms.

In the current study, we found a differential effect of upadacitinib compared to dupilumab on the methylome profile of the skin-homing T helper cells. Dupilumab reversed 4 out of 6 AD-related DMRs toward a methylome profile more closely resembling HC. Among these, TSPAN32 is reported to play an immunoregulatory role in T helper cells and is downregulated upon T cell activation in memory T cells. Furthermore, lower levels of *Tetraspanin 32* (TSPAN32) expression in PBMCs is associated with multiple sclerosis relapse [32]. Additionally, *interferon-stimulated positive regulator* (BISPR) is a long non-coding RNA, closely associated with *Tetherin* (BST2). Both are induced in response to interferon-α (IFNα) [33, 34] and play a role in T cell antiviral response [35]. *BST2* is highly expressed in lesional AD skin and acts as a potent itch mediator that functions downstream of IL27-mediated pruritus response [36]. These observations suggest that dupilumab may partially restore immune homeostasis in AD by epigenetically modulating key regulators of T cell activation, antiviral responses, and pruritus. Alternatively, upadacitinib enhanced the majority of AD-related DMRs, further shifting them towards the methylome profile characteristic of AD. Additionally, upadacitinib had much broader effect on the DNA methylome, inducing more AD-unrelated DMRs compared to dupilumab. It is noteworthy that our study findings align with clinical observations in patients treated with dupilumab and upadacitinib. JAK inhibitors, including upadacitinib, have been reported to cause a wider range of adverse events compared to dupilumab. Additionally, treatment discontinuation of JAK inhibitors is associated with a rapid disease flare [8], whereas dupilumab is associated with longer periods of remission upon treatment cessation [5, 7]. Consistent with this, we previously reported persistently elevated serum TARC levels in AD patients treated with a JAK inhibitor, despite a good clinical response [37]. These findings highlight the contrasting epigenetic footprints of the two treatments, with dupilumab partially restoring an AD-associated methylation signature toward a healthy state, while upadacitinib exerts broader and more complex effects that may reinforce or diversify the AD-related epigenetic landscape. It is important to note that the dupilumab-treated group was biologic treatment–naïve, unlike the upadacitinib group, in which the majority of patients had been previously treated with either dupilumab or other JAK inhibitors. However, no significant differences in the methylation profiles were observed prior to the initiation of dupilumab or JAK inhibitor therapy, indicating that baseline epigenetic signatures were comparable between the groups.

Our analysis further highlighted epigenetic regulation of IL13 and IL22 protein expression at the DNA methylation level, whereas no such regulation was observed for IL4 and IL5. In line with this, a previous report demonstrated no significant change in the *IL4* and *IL5* gene methylation in blood of the allergic patients compared to healthy donors [38]. Interestingly, the *IL13* promoter showed further hypomethylation following upadacitinib treatment. In contrast, dupilumab treatment increased *IL13* promoter methylation, shifting it toward HC phenotype. This indicates that among all Th2 cytokines, only IL13 is epigenetically modulated by both treatments in opposite directions. This finding may have implications for disease remission or flare given the key role of IL13 in AD pathogenesis. The lack of reduction in IL13 protein levels in our cohort is mostly derived by one patient outlier and our small sample size. However, in a larger cohort, we previously demonstrated that dupilumab treatment decreases IL13-expressing skin-homing T helper cells after four weeks of treatment, with levels remaining stably low for up to 52 weeks [39].

In conclusion, this study is the first to show that dupilumab and upadacitinib exert distinct, cell-specific effects on the DNA methylome in AD. Despite the small sample size, we identified differential epigenetic responses in skin-homing T helper cells, with upadacitinib inducing broader, but predominantly AD-like methylation changes, while dupilumab partially corrected disease-associated patterns. These findings align with clinical observations, supporting a favorable safety profile and potential for treatment tapering with dupilumab. Our results underscore the relevance of epigenetic profiling in AD and highlight the need for future studies exploring other immune cell types, additional epigenetic layers, the effects of prolonged treatment, and the durability of changes following treatment discontinuation.

## Abbreviations

AD: atopic dermatitis
HC: non-atopic healthy controls
DMRs: differentially methylated regions
DMPs: differentially methylated positions
DEGs: differentially expressed genes

## Data Availability Statement

The data that support the findings of this study are available from the corresponding author upon reasonable request.

## Acknowledgments

We would like to thank our patients included in this manuscript for participating in the BioDay registry. The BioDay registry is supported by Eli Lilly and Company, Sanofi, Leo Pharma and Pfizer. We also thank Health~Holland, the Top Sector Life Sciences & Health, for providing financial support for the researchers (A.E).

