## Supplementary Figure S1 for "Impact of dupilumab and upadacitinib treatment on epigenetic and transcriptomic alterations in skin-homing T cells of atopic dermatitis patients"

**Supplementary Appendix**

**Supplementary Figures**

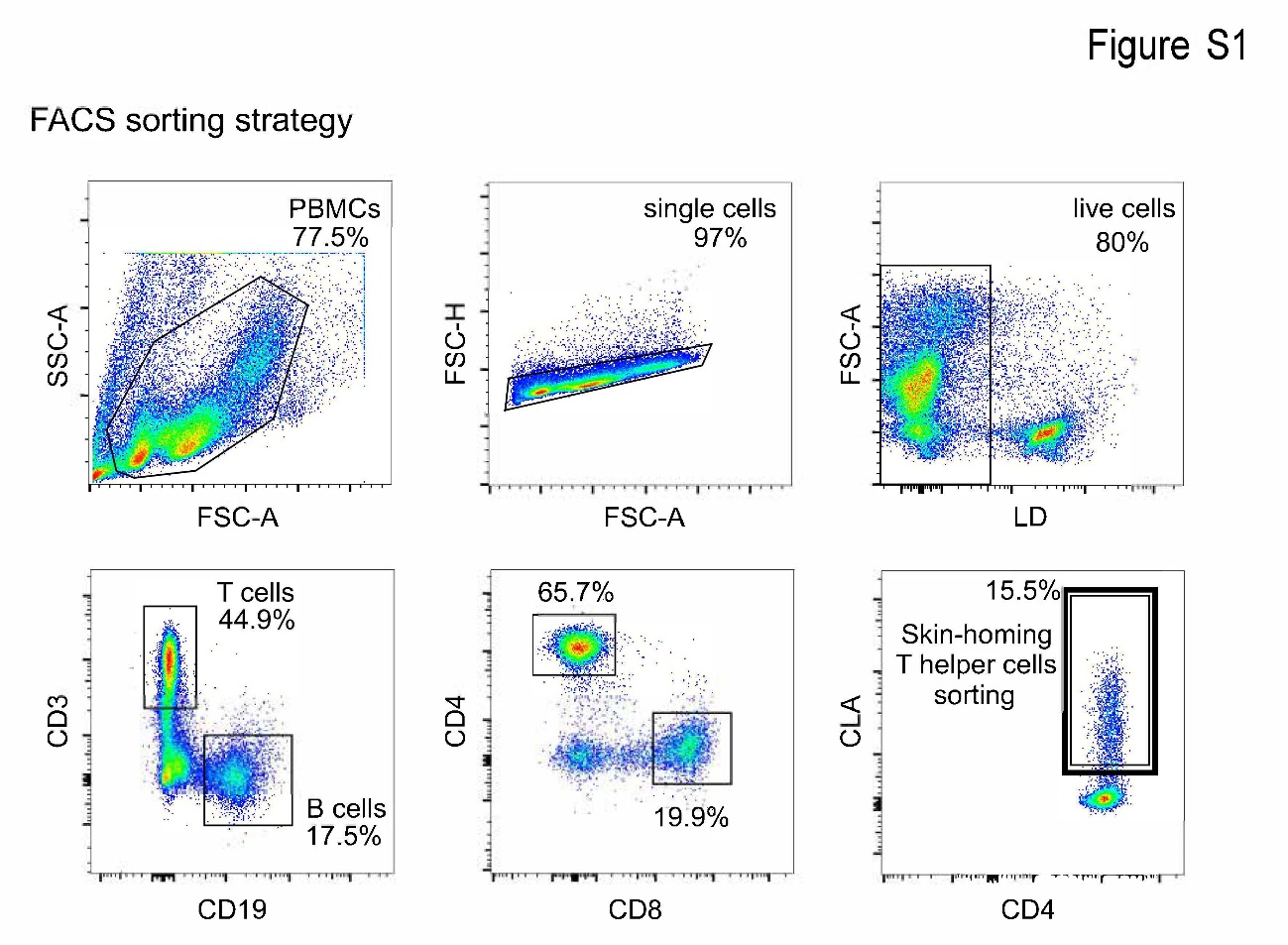

**Figure S1. Flow cytometry gating strategy for skin-homing T helper cells sorting.**

FACS plots show gating strategy for sorting CD3^+^CD4^+^CLA^+^ T cells.
