## Supplementary Table S1 for "Impact of dupilumab and upadacitinib treatment on epigenetic and transcriptomic alterations in skin-homing T cells of atopic dermatitis patients"

**Supplementary Appendix**

**Supplementary Tables**

**Table S1. Flowcytometry antibodies.**

| **Target** | | **Fluorochrome** | **Company** | **Catalogue #** | **Dilution** |
| --- | --- | --- | --- | --- | --- |
|  | Sorting antibodies | | | | |
| - Fixable viability dye | | eF780 | Thermo Fisher Scientific | 65-0865-18 | 1:1000 |
| - CD3 | | BV605 | Biolegend | 300460 | 1:100 |
| - CD4 | | BV785 | Biolegend | 300554 | 1:50 |
| - CD8 | | PE-Cy7 | BD | 335822 | 1:200 |
| - CLA | | Pacific blue | Biolegend | 321308 | 1:200 |
| Flowcytometry analysis antibodies | | | |  |  |
| - Fixable viability dye | | eF506 | Thermo Fisher Scientific | 65-0866-18 | 1:1000 |
| - CD3 | | BV605 | Biolegend | 300460 | 1:100 |
| - CD4 | | BV785 | Biolegend | 300554 | 1:50 |
| - CD8 | | APC-Cy7 | BD | 557834 | 1:50 |
| - CLA | | Pacific blue | Biolegend | 321308 | 1:200 |
| - Ki67 | | FITC | DAKO | F7268 | 1:200 |
| - IL4 | | BV711 | BD | 564112 | 1:12.5 |
| - IL5 | | PE | Biolegend | 500904 | 1:50 |
| - IL13 | | Percp.Cy5.5 | Biolegend | 501912 | 1:50 |
| - IL22 | | APC | eBioscience | 17-7222-82 | 1:40 |
