## Supplementary Table S2 for "Impact of dupilumab and upadacitinib treatment on epigenetic and transcriptomic alterations in skin-homing T cells of atopic dermatitis patients"

**Supplementary Appendix**

**Supplementary Tables**

**Table S2. List of genes annotated to DMRs modulated by dupilumab (p<0.1).**

| **Gene name** | **Methylation MeanFC** | **Methylation maxDiff** |
| --- | --- | --- |
| **AD-related DMRs changes** | | |
| BST2 | -1,85 | -0,429196177 |
| MTMR1 | -1,81 | -0,446556459 |
| BISPR | -1,80 | -0,429196177 |
| TSPAN32 | -1,70 | -0,504773008 |
| SLC1A5 | -1,56 | -0,56677446 |
| ALPK2 | 2,18 | 0,383776123 |
| **Newly induced DMRs changes** | | |
| CXCR6 | -2,83 | -0,550469637 |
| VPS41 | -2,69 | -0,470018019 |
| CXCR3 | -1,82 | -0,404424058 |
| DTX3L | -1,77 | -0,319070043 |
| LETM2 | -1,63 | -0,79636842 |
| MVB12A | -1,54 | -0,429196177 |
| IL32 | -1,43 | -0,382085097 |
| SEPTIN9 | -1,39 | -0,648752531 |
| SEMA4A | -1,37 | -0,336042937 |
| NR2F1-AS1 | 1,37 | 0,34635864 |
| TACR1 | 1,41 | 0,459108478 |
| SPAG6 | 1,67 | 0,305299816 |
| FLRT2 | 1,70 | 0,450215611 |
