## Supplementary Table S3 for "Impact of dupilumab and upadacitinib treatment on epigenetic and transcriptomic alterations in skin-homing T cells of atopic dermatitis patients"

**Supplementary Appendix**

**Supplementary Tables**

**Table S3. List of genes annotated to DMRs modulated by upadacitinib (p<0.1).**

| **Gene name** | **Methylation MeanFC** | **Methylation maxDiff** |
| --- | --- | --- |
| **AD-related DMRs changes** | | |
| IL21 | -6,09 | -1,123972878 |
| TNFAIP8 | -2,69 | -1,074784367 |
| CFLAR | -1,99 | -0,755828959 |
| PDCD1 | -2,72 | -0,73815234 |
| LOC283387 | -2,30 | -0,736994782 |
| TGIF1 | -1,91 | -0,666145194 |
| VOPP1 | -2,04 | -0,616453231 |
| PLA1A | -2,80 | -0,59478673 |
| ANXA6 | -1,89 | -0,541668052 |
| BRWD1 | -2,39 | -0,539990947 |
| BRWD1-AS2 | -2,39 | -0,539990947 |
| ATG4A | -2,22 | -0,533138081 |
| TIGIT | -2,46 | -0,523288105 |
| GPR137 | -1,77 | -0,522515194 |
| TNFSF4 | -2,65 | -0,521719312 |
| OTUD5 | -1,39 | -0,495724897 |
| IL13 | -1,65 | -0,490357457 |
| PLEC | -1,39 | -0,468052198 |
| GNAS | -1,11 | -0,456323372 |
| MAP1LC3B2 | -1,77 | -0,443343908 |
| MIR1207 | -1,70 | -0,424207055 |
| ALPK2 | -2,08 | -0,415329158 |
| CAPN2 | -1,97 | -0,4115426 |
| LOC100507156 | -2,20 | -0,4097018 |
| PDCD4-AS1 | -2,27 | -0,399428343 |
| SYNGAP1 | -2,06 | -0,396250688 |
| HSD11B1-AS1 | -2,02 | -0,389144166 |
| C1RL-AS1 | -1,84 | -0,388843776 |
| C18orf21 | -2,54 | -0,388692533 |
| LGALS1 | -2,30 | -0,346566333 |
| ADAMTS16 | -1,65 | -0,345643525 |
| F5 | -1,95 | -0,327688327 |
| BCL9L | -1,82 | -0,320209003 |
| TNFRSF4 | -2,23 | -0,257565149 |
| SULF2 | -1,28 | -0,2507941 |
| MSL3 | -1,55 | -0,249991346 |
| LTB | 2,60 | 0,415122346 |
| FGF1 | 1,86 | 0,425214662 |
| PTHLH | 2,42 | 0,425244699 |
| FOSL2-AS1 | 2,71 | 0,715918239 |
| **Newly induced DMRs changes** | | |
| SMARCA2 | -3,55 | -1,009398218 |
| FAIM | -2,40 | -0,736154796 |
| PDE3A | -2,44 | -0,715676637 |
| CXCR5 | -2,96 | -0,702118178 |
| TAF6 | -2,85 | -0,694465605 |
| NUDC | -2,99 | -0,686749528 |
| MIR12129 | -1,48 | -0,679724842 |
| CD300LG | -3,34 | -0,672907053 |
| SPATA6L | -1,84 | -0,670766272 |
| CD109-AS1 | -2,30 | -0,668326093 |
| TMEM182 | -1,82 | -0,651619575 |
| SPOCK2 | -3,47 | -0,646051223 |
| LMO1 | -2,17 | -0,644782631 |
| MDFIC | -1,91 | -0,624935705 |
| TRPA1 | -1,88 | -0,619437984 |
| FSBP | -1,58 | -0,602988584 |
| KCNIP4 | -1,75 | -0,602443577 |
| EGR2 | -2,79 | -0,598193862 |
| IDH2 | -2,48 | -0,590590219 |
| IDH2-DT | -2,67 | -0,590590219 |
| EFCAB10 | -3,02 | -0,583705467 |
| ELMO1 | -1,47 | -0,572138745 |
| CACNA1G | -1,50 | -0,567720033 |
| NCOA7 | -2,65 | -0,556358562 |
| DMAC2L | -1,53 | -0,545164671 |
| L2HGDH | -1,32 | -0,545164671 |
| PDE4A | -1,52 | -0,543816557 |
| CDK5R2 | -1,60 | -0,54326511 |
| VSTM2A | -1,67 | -0,535412324 |
| IKBKB | -2,90 | -0,534518356 |
| IKBKB-DT | -2,97 | -0,534518356 |
| DPYSL4 | -1,41 | -0,533143333 |
| HIC1 | -1,42 | -0,532969855 |
| GPR34 | -2,94 | -0,522955577 |
| ZBED9 | -1,33 | -0,515281664 |
| TBC1D12 | -2,60 | -0,509319325 |
| TMEM216 | -1,94 | -0,507323981 |
| ZNF211 | -1,49 | -0,504010793 |
| NTRK2 | -2,00 | -0,502557344 |
| ATOH1 | -2,03 | -0,498750452 |
| ZER1 | -2,23 | -0,491849219 |
| MMP9 | -3,42 | -0,487941506 |
| TM2D1 | -2,07 | -0,485057821 |
| H1-1 | -2,14 | -0,482825374 |
| RUFY3 | -1,35 | -0,48140713 |
| AIMP1 | -1,78 | -0,480507772 |
| SNX10-AS1 | -2,32 | -0,478785557 |
| INGX | -1,82 | -0,477949189 |
| TCF7 | -1,82 | -0,476245059 |
| DRD1 | -2,28 | -0,468494805 |
| EDARADD | -2,50 | -0,467461948 |
| PLK2 | -2,13 | -0,46699916 |
| MGAT4A | -2,10 | -0,464671769 |
| TRIM36 | -1,68 | -0,464587836 |
| RAB1B | -1,96 | -0,464377039 |
| CAV1 | -1,98 | -0,458109895 |
| HLTF | -2,00 | -0,457998545 |
| LOC101927932 | -1,26 | -0,456323372 |
| FAM169A | -1,93 | -0,451540632 |
| LOC441086 | -1,93 | -0,451540632 |
| C2CD2L | -2,76 | -0,450558246 |
| LINC00852 | -2,42 | -0,449983135 |
| ZNF706 | -2,35 | -0,443966759 |
| TRAPPC13 | -1,54 | -0,440787166 |
| GANC | -1,44 | -0,440745471 |
| IMPG2 | -1,39 | -0,437356981 |
| RUBCNL | -1,76 | -0,436717351 |
| SLC22A15 | -1,34 | -0,435444562 |
| MPRIP | -2,25 | -0,432916392 |
| ADAMTS5 | -1,38 | -0,427779773 |
| AK5 | -2,19 | -0,427523317 |
| TSPAN13 | -2,73 | -0,427512909 |
| NUAK1 | -2,03 | -0,42654339 |
| MIB2 | -2,32 | -0,424288377 |
| PEG10 | -1,45 | -0,423834948 |
| RAB27A | -2,03 | -0,420866483 |
| DELE1 | -2,07 | -0,418022192 |
| PPAN | -1,97 | -0,4171899 |
| FOXC1 | -1,88 | -0,416934863 |
| HCG18 | -2,11 | -0,413683662 |
| RAI2 | -2,47 | -0,413388306 |
| LINC00473 | -1,87 | -0,412056424 |
| IGF2 | -1,37 | -0,408220024 |
| PDZRN3 | -1,34 | -0,408177345 |
| PSMD10 | -1,98 | -0,404490402 |
| SLC38A11 | -1,38 | -0,404079514 |
| ZNF607 | -1,62 | -0,403851059 |
| GATA4 | -1,37 | -0,40227929 |
| ARHGAP20 | -1,73 | -0,399792943 |
| DPY19L2P4 | -1,50 | -0,398637516 |
| RTN4 | -1,40 | -0,397810173 |
| PIGV | -1,89 | -0,396524131 |
| CUTA | -1,54 | -0,396250688 |
| LRIG3 | -1,76 | -0,395801582 |
| KIFC1 | -0,99 | -0,395575813 |
| CD109 | -2,02 | -0,395221227 |
| RUNDC3B | -1,88 | -0,394839424 |
| RRM1 | -1,46 | -0,393794024 |
| SOX2 | -1,48 | -0,391823586 |
| ATP5MC2 | -1,23 | -0,390942901 |
| ETV5 | -1,79 | -0,390570975 |
| GFRA1 | -1,89 | -0,389948561 |
| ZNF516 | -1,61 | -0,388876621 |
| IFIT2 | -1,91 | -0,388499172 |
| NIM1K | -1,81 | -0,38641003 |
| ZNF256 | -1,61 | -0,385932499 |
| RABGEF1 | -1,47 | -0,384387098 |
| WFDC3 | -1,92 | -0,383866025 |
| SSX2IP | -1,83 | -0,382125257 |
| STK26 | -1,53 | -0,376876526 |
| DTX1 | -1,77 | -0,373926242 |
| IRS2 | -1,26 | -0,371503684 |
| ASIC2 | -1,77 | -0,367053453 |
| HOATZ | -1,93 | -0,366373737 |
| SMAD5 | -1,98 | -0,364461916 |
| LINC01535 | -2,19 | -0,363344986 |
| MECP2 | -2,40 | -0,362013868 |
| GBE1 | -1,50 | -0,361628115 |
| TTC14 | -1,82 | -0,361309378 |
| MINPP1 | -1,51 | -0,360952313 |
| IFNGR2 | -2,06 | -0,360794707 |
| DBF4 | -2,25 | -0,360617165 |
| PLGRKT | -1,79 | -0,360219684 |
| PDE5A | -1,51 | -0,359742035 |
| VCAN | -2,14 | -0,3586859 |
| SHISA9 | -1,68 | -0,357913976 |
| ETV1 | -2,11 | -0,354888528 |
| LUZP1 | -1,74 | -0,353490953 |
| H3-3A | -1,44 | -0,351404322 |
| MID1IP1 | -1,44 | -0,347330545 |
| TBC1D2B | -1,65 | -0,343763211 |
| GOLGA3 | -2,72 | -0,34229385 |
| LUZP2 | -1,54 | -0,341491308 |
| RRAGD | -2,04 | -0,339781728 |
| MCMBP | -1,83 | -0,337292116 |
| PACRG | -1,57 | -0,335661856 |
| KCNAB2 | -1,68 | -0,334776941 |
| RETREG1-AS1 | -2,14 | -0,333110408 |
| SOX11 | -1,61 | -0,327074123 |
| ALX1 | -2,15 | -0,324776682 |
| ZCCHC24 | -1,45 | -0,324232593 |
| TRIM16 | -2,31 | -0,32096198 |
| FUCA1 | -1,63 | -0,320235573 |
| PEX2 | -2,13 | -0,313803621 |
| GFRA2 | -1,44 | -0,312052613 |
| ZNF345 | -1,55 | -0,3119896 |
| ASB5 | -1,71 | -0,310909431 |
| SLC29A1 | -2,05 | -0,310286202 |
| WNT2 | -1,77 | -0,309030741 |
| BICD2 | -2,11 | -0,308924451 |
| PHTF2 | -1,60 | -0,306042061 |
| OPHN1 | -1,89 | -0,304414536 |
| RECQL | -1,34 | -0,300926533 |
| PFKFB2 | -2,05 | -0,300761613 |
| SNX22 | -1,78 | -0,300004198 |
| NCMAP | -1,82 | -0,299092472 |
| LINC01503 | -1,86 | -0,298308159 |
| CTSD | -2,03 | -0,296929341 |
| TP53TG5 | -2,05 | -0,29522113 |
| NBL1 | -1,66 | -0,292505492 |
| ZNF621 | -1,63 | -0,291813357 |
| DECR2 | -1,77 | -0,291237203 |
| LINC02585 | -1,62 | -0,290541572 |
| HPCA | -1,66 | -0,288833346 |
| LOC100272217 | -1,55 | -0,28875667 |
| SMC3 | -1,60 | -0,28728817 |
| ZNF134 | -1,74 | -0,285430072 |
| CCDC34 | -1,47 | -0,284384734 |
| MFSD9 | -1,91 | -0,283276911 |
| SLCO5A1 | -1,38 | -0,282639868 |
| TMEM268 | -1,94 | -0,282620932 |
| YARS1 | -1,73 | -0,278036874 |
| LMBRD2 | -1,75 | -0,27661209 |
| SKP2 | -1,68 | -0,27661209 |
| STAC | -1,72 | -0,276601932 |
| HTR1B | -1,32 | -0,27397276 |
| MID1IP1-AS1 | -1,57 | -0,271998861 |
| IDS | -1,64 | -0,267674471 |
| HCFC1 | -1,64 | -0,266575616 |
| AIP | -1,84 | -0,265826155 |
| EYA4 | -1,40 | -0,262243083 |
| DACT1 | -1,62 | -0,252166713 |
| RNF130 | -1,64 | -0,248523565 |
| ABCC9 | -1,54 | -0,248087072 |
| PPP1R7 | -1,72 | -0,240669378 |
| TMEM256 | -1,33 | -0,235288505 |
| TMEM256-PLSCR3 | -1,28 | -0,235288505 |
| MRM2 | -1,15 | -0,230131212 |
| KCNK12 | -1,26 | -0,223744808 |
| SLC25A14 | -1,30 | -0,221160021 |
| HELZ2 | -1,53 | -0,22030468 |
| TCEAL2 | -1,60 | -0,209637703 |
| FSCN1 | -1,14 | -0,192186744 |
| PPM1J | -1,49 | -0,188978546 |
| TNNI1 | 1,72 | 0,227906846 |
| MYBPHL | 1,86 | 0,257682331 |
| LINC02751 | 1,73 | 0,282849342 |
| LOC101928438 | 2,04 | 0,284959646 |
| LOC101928881 | 1,80 | 0,292980022 |
| GPHA2 | 1,94 | 0,295853971 |
| OR2B2 | 1,90 | 0,297812356 |
| KLK9 | 2,38 | 0,300218935 |
| CAPN13 | 2,17 | 0,301730323 |
| MIR26A2 | 2,53 | 0,308507777 |
| FCRL6 | 1,75 | 0,327383264 |
| BRD4 | 1,91 | 0,328079525 |
| OPALIN | 2,55 | 0,338210076 |
| C15orf32 | 1,94 | 0,340663183 |
| ACTBL2 | 2,28 | 0,343520661 |
| SLC16A3 | 1,55 | 0,34934127 |
| TRPM1 | 1,22 | 0,349714272 |
| TLR6 | 2,00 | 0,378386346 |
| PPP1R1C | 2,19 | 0,384255121 |
| PTAFR | 2,60 | 0,390987621 |
| LINC02387 | 2,34 | 0,393750218 |
| FAM216B | 2,49 | 0,398986788 |
| LINC00839 | 2,61 | 0,404961398 |
| LINC00636 | 2,98 | 0,405125879 |
| LOC101926934 | 2,18 | 0,408425302 |
| TDRD6 | 1,97 | 0,408425302 |
| LOC101929154 | 2,16 | 0,413443614 |
| KALRN | 1,56 | 0,415836728 |
| CCR6 | 2,15 | 0,421057234 |
| CCDC70 | 2,39 | 0,423055727 |
| MZB1 | 2,59 | 0,429143844 |
| ANKRD22 | 3,21 | 0,438353785 |
| IKZF4 | 2,49 | 0,441634567 |
| OLFML1 | 2,22 | 0,446690082 |
| SLC36A3 | 2,58 | 0,45985514 |
| CCR5AS | 2,79 | 0,461413376 |
| CCRL2 | 2,56 | 0,461413376 |
| IL1RN | 2,63 | 0,471262375 |
| XPNPEP2 | 2,71 | 0,487132443 |
| SOAT2 | 2,28 | 0,501526828 |
| CD27 | 2,66 | 0,53506013 |
| OR51S1 | 3,45 | 0,55130891 |
| CXCR3 | 3,63 | 0,572621322 |
| STRA6 | 2,64 | 0,578389067 |
| SCARNA27 | 3,57 | 0,592561381 |
| SNORA80B | 3,74 | 0,656200085 |
| IFIT3 | 2,62 | 0,678799904 |
